# Genome-wide prediction of synthetic rescue mediators of resistance to targeted and immunotherapy

**DOI:** 10.1101/284240

**Authors:** Avinash Das, Joo Sang Lee, Gao Zhang, Zhiyong Wang, Ramiro Iglesias-Bartolome, Tian Tian, Zhi Wei, Benchun Miao, Nishanth Ulhas Nair, Olga Ponomarova, Adam A. Friedman, Arnaud Amzallag, Tabea Moll, Gyulnara Kasumova, Patricia Greninger, Regina K. Egan, Leah J. Damon, Dennie T. Frederick, Allon Wagner, Kuoyuan Cheng, Seung Gu Park, Welles Robinson, Kevin Gardner, Genevieve Boland, Sridhar Hannenhalli, Meenhard Herlyn, Cyril Benes, J. Silvio Gutkind, Keith Flaherty, Eytan Ruppin

## Abstract

Most patients with advanced cancer eventually acquire resistance to targeted therapies, spurring extensive efforts to identify molecular events mediating therapy resistance. Many of these events involve *synthetic rescue (SR) interac*tions, where the reduction in cancer cell viability caused by targeted gene inactivation is rescued by an adaptive alteration of another gene (the *rescuer*). Here we perform a genome-wide prediction of SR rescuer genes by analyzing tumor transcriptomics and survival data of 10,000 TCGA cancer patients. Predicted SR interactions are validated in new experimental screens. We show that SR interactions can successfully predict cancer patients’ response and emerging resistance. Inhibiting predicted rescuer genes sensitizes resistant cancer cells to therapies synergistically, providing initial leads for developing combinatorial approaches to overcome resistance proactively. Finally, we show that the SR analysis of melanoma patients successfully identifies known mediators of resistance to immunotherapy and predicts novel rescuers.

## INTRODUCTION

Despite major advances in cancer therapies, many patients eventually succumb to emerging resistance. Recent experimental and clinical studies have successfully characterized tumor-specific molecular signatures of resistance to targeted therapies through DNA and RNA sequencing(*1-8*). However, these studies require the arduous collection and molecular profiling of paired pre- and post-treatment tumor biopsies(*9*) and cannot be conducted for drugs at early stages of their development. Thus, the development of a computational approach that can expedite the identification of resistance determinants from existing large-scale cancer cohorts is warranted.

We set out to predict *synthetic rescue (SR)* interactions (*1-3, 10, 11*), which are a generalization of suppressor interactions(*12*). Suppressor interactions, recently identified in yeast genome-wide(*11*), denote a functional interaction where following the inactivation of specific genes cells suppress additional genes to escape from harmful alterations(*13-15*). Their generalization, SR interactions, denotes a functional interaction where a fitness reducing alteration due to inactivation of one gene (termed the *vulnerable* gene) is compensated by altered activity (down-regulation or up-regulation) of another, *rescuer* gene(*9, 16-18*) (Figure 1a). As rescue events are required to compensate for fitness reducing alterations occurring during the natural evolution of cancer (*19*), one may expect to detect the SR interactions forged in evolving tumors, even untreated ones(*20-22*). When a *vulnerable* gene is targeted by an anti-cancer drug(*23*), such SR interactions may manifest by changes in the activity of its interacting *rescuer* gene(s), thus mediating drug resistance. Both primary and adaptive resistance could be mediated by SR mechanisms.

**Figure 1.**
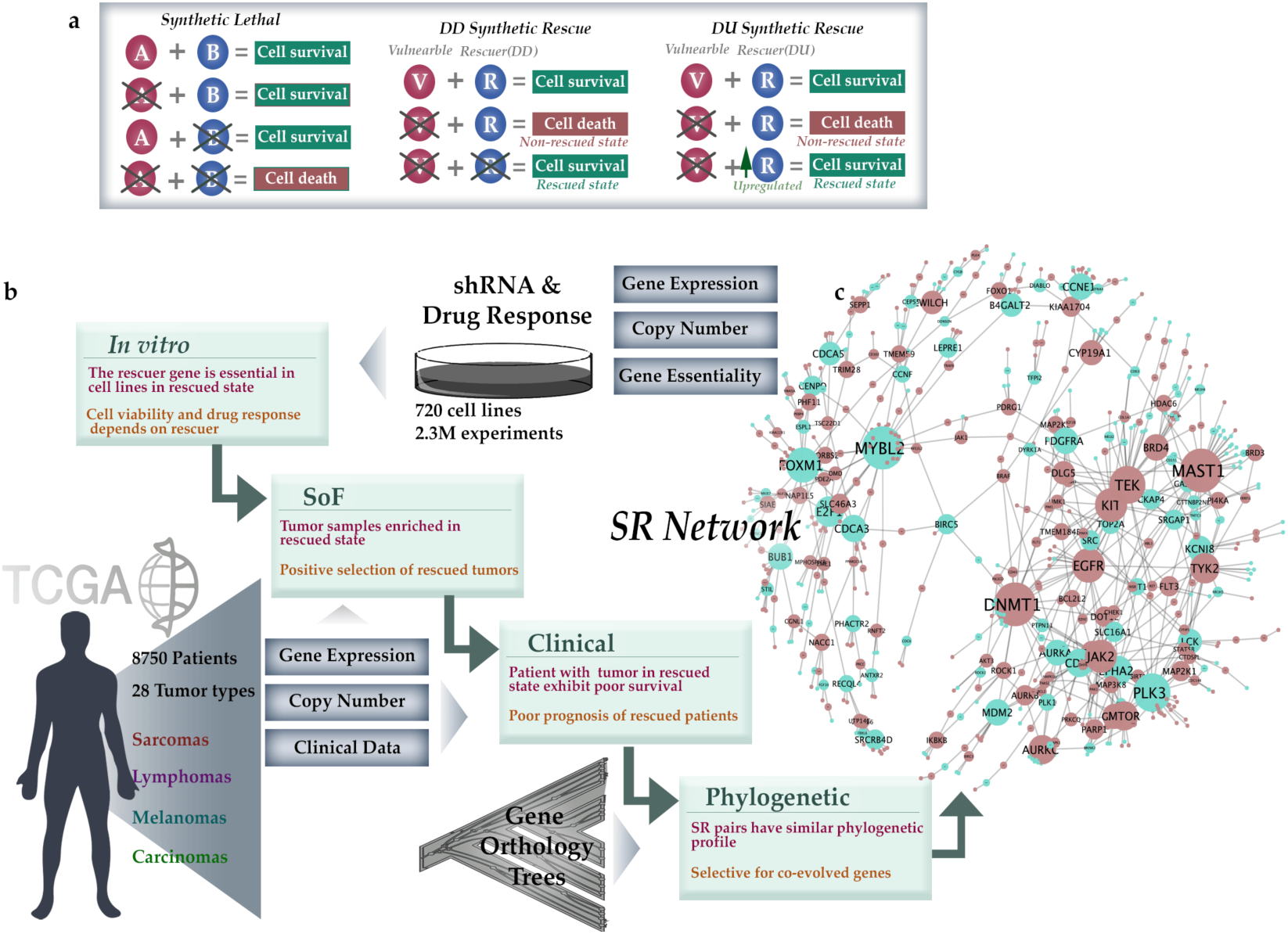
The INCISOR pipeline and the resulting SR network: **(a)** The phenotypic effects of altering interacting gene partners in SL, DD-SR, and DU-SR interactions. **(b)** The **four inference steps of INCISOR** and the datasets analyzed (Methods, SoF stands for the survival of the fittest). The SR property tested (in red) and rationale (in brown) of each step are also displayed. **(c)** The resulting DU-SR network (purple nodes denote vulnerable genes and green rescuer genes; the size of nodes is proportional to the number of interactions they have). The complete network is provided in Fig. S1f.

## RESULTS

We and others have recently developed computational approaches to identify synthetic lethal (SL) interactions(*24-30*), a widely studied class of genetic interactions. While SL interactions pinpoint molecular vulnerabilities in tumors that can be exploited to target them(*31, 32*), SR interactions can rescue the cells from such vulnerabilities by actively modifying the interacting rescuers, leading to therapy resistance. As drugs mainly inhibit target genes, we focus herewith on two types of SR interactions (**Figure 1a**); (1) **DU**-SR interactions, where the **D**ownregulation of a vulnerable gene is rescued by **the U**pregulation of a rescuer gene (*2-4, 33-36*); and (2) **DD**-SR (suppressor) interactions, where the **D**ownregulation of a vulnerable gene is rescued by the **D**ownregulation of a rescuer gene (*10, 11, 19, 37, 38*).

### The INCISOR Pipeline and the resulting Cancer SR networks

To predict SR interactions, we developed a new data mining approach, termed “IdeNtification of ClinIcal Synthetic Rescues in cancer” (INCISOR). Conceptually, INCISOR simply combines multiple lines of evidence – experimental, tumor transcriptomics, clinical and gene phylogeny – to ascertain if a gene pair is likely to be SR. Here we describe the algorithm of INCISOR that predicts DU-SR interactions, where the rescue event is mediated by over-expression (DD-SR prediction follows an analogous approach, Methods, Suppl. Information 2, Fig. S1g). INCISOR analyzes *in vitro* screens and evaluates the extent which gene phylogeny, molecular, and survival data of patient tumor support the screens. It selects the clinically-relevant SR pairs that are supported by all four line of evidence outlined below. The specific order in which the following four steps are applied sequentially in INCISOR was chosen to minimize the computational cost (**Figure 1b**, see Methods for details):

(1) *In vitro* essentiality screens: This step mines *in vitro* genome-wide shRNA(*39-42*) and drug response screens(*43, 44*) composed of 2.3 million measurements in 720 cancer cell lines. Adopting a recent approach(*29*), INCISOR systematically searches for candidate SR pairs. In cell lines with a given gene knockdown, it searches the cell-line’s transcriptome for genes whose up-regulation is associated with increased cell growth. We term the first gene a vulnerable (V) gene and the second a candidate (DU) rescuer (R) gene.
(2) Molecular survival of the fittest (SoF): By analyzing TCGA gene expression and somatic copy number alterations (SCNA) of 8,749 patients across 28 cancer types, INCISOR selects candidate SR pairs from step 1 that are observed in their *rescued* state (gene R is specifically upregulated when gene V is inactive) significantly more than expected. This enrichment testifies to positive selection of samples in the rescued state, a key property of SR interactions.
(3) Patient survival screening: This step further selects those candidate SRs whose *rescued* state in TCGA tumor samples exhibits worse patient’s survival, as the reduced survival can serve as an indicator of increased tumor fitness. INCISOR uses a stratified Cox proportional hazard model to establish this relation while controlling for confounding factors including cancer type, sex, age, genomic instability, tumor purity(*45*), and ethnicity.
(4) Phylogenetic screening: Because functionally interacting genes known to co-evolve(*28*) in a species, we select SR pairs composed of genes with high phylogenetic similarity. The top 5% of phylogenetically similar pairs among the ones passing the previous steps are chosen as the final set of putative SR pairs.

The resulting *DU-SR* network, which is composed of all the pairwise interactions that pass all four steps described above, is scale-free (**Figure 1c**, Table S2,3) and consists of 1259 genes and 1195 interactions (see Suppl. Information 2.1 for DD-SR; interactive networks available online, Methods; Table S4,5). Gene enrichment analysis revealed that the network nodes are enriched in cancer and resistance pathways (Suppl. Information 3.5-3.7). We also find that the activation of predicted rescuers increases with advanced cancer stages (Suppl. Information 3.9, Fig. S2g,h).

### Benchmarking INCISOR against a collection of published DU-SR Interactions

We first benchmarked the *DU-SR* predictions via a comparison to genes whose over-expression rescues cancer cells, via a set of genes that were previously shown to mediate cancer drug resistance(*2, 36, 46-52*) (Table S9, Methods, Suppl. Information 4.1). INCISOR successfully identified these published rescuer genes with AUCs of 70-85% (mean precision of 46% at 50% recall; Fig. S3o, Methods, Suppl. Information 4.1). We additionally tested and successfully validated predicted SR interactions using published data of patient-derived *in vitro* (*53*) and mouse xenograft models (*54*) (Suppl. Information 4.4, 4.5). As large cohorts of published rescue interactions are still quite scarce, we turned to conduct four new *in vitro* experiments to further test emerging rescue predictions of INCISOR of interest.

### Experimental testing of predicted DD-SR interactions of mTOR in Head & Neck Cancer cell lines

Our first experiment tested *DD-SR* interactions involving mTOR, a key growth regulating kinase in Head & Neck Cancer. To test the predicted rescue interactions involving mTOR, we knocked down (KD) 2200 genes in an experimental screen in a head and neck cancer cell line (HN12) and experimentally identified the (DD) rescue events occurring due to a subsequent mTOR inhibition by rapamycin treatment. 45 of these KDs, about 2.1%, were rescued by mTOR inhibition in the screen (Table S10, Methods). Independently, we applied INCISOR to identify genes that are predicted to be rescued by mTOR inhibition in a statistically significant manner (FDR = 0.05,). INCISOR predicted 17 such *DD* rescuer genes (Methods), 11 of which indeed overlapped with the 45 interactions identified experimentally (Fig. S5b). This yields a precision level of ~65% and recall of ~25% (**Figure 2a, false positive rate < 0.003**), a 31-fold increase over the 2.1% precision expected by chance. The validated rescuers were enriched with transcription factors, FoxO signaling and stress response genes (Table S30). We further validated the predicted DD-SR interactions of mTOR via multiple published *in vitro* shRNA(*39-42*) and drug response screens(*43, 44*) (Suppl. Information 4.2, 4.3).

**Figure 2:**
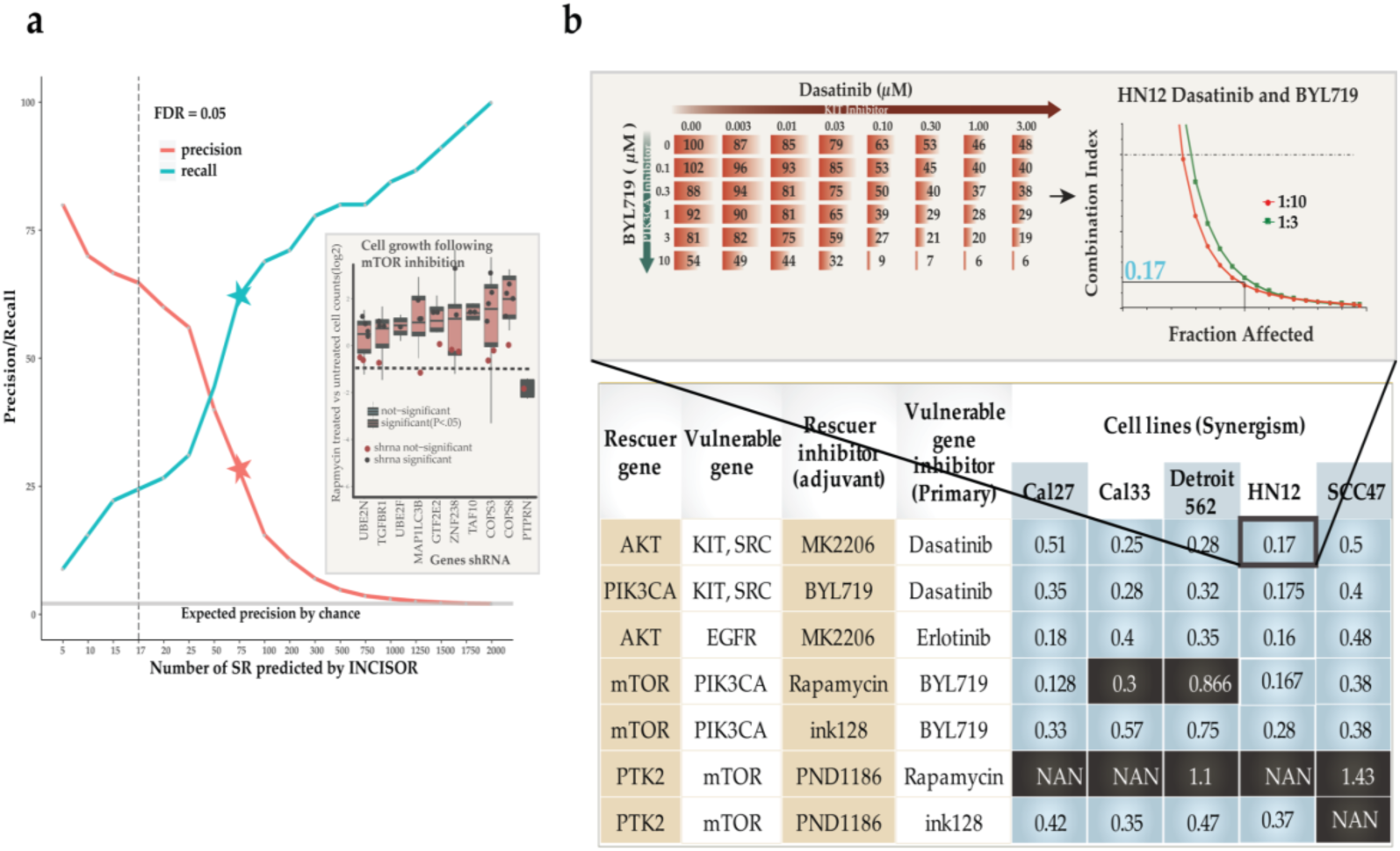
Large-scale *in vitro* experiments testing predicted SR interactions in Head and neck cancer (a) Evaluation of predicted SR (DD) interactions in a large-scale shRNA H&N HN12 cell-line screen. The Y-axis displays the precision and recall of INCISOR-predicted SRs in identifying the 45 experimentally determined *SR-DD* rescuers of mTOR. The vertical dashed line denotes a threshold of FDR = 0.05 over the predicted INCISOR interaction scores. The stars indicate precision and recall at a threshold level where INCISOR identifies 75 genes as *SR-DD* rescuers. The horizontal line (in gray) shows the precision expected by the random chance. The inset displays top 10 predicted genes whose knockdowns are rescued by mTOR inhibition. **(b) Experimental validation of predicted synergistic SR-based combinational therapies in head and neck cancer**: A table summarizing the experimentally observed synergism between primary drugs and their predicted rescuer-targeting treatments in 5 HNSC cell lines, based on drug treatment experiments. Synergism was estimated using a standard Fa-CI analysis. The table displays the average combination index (CI; synergism CI < 1, additivity effect CI = 1, antagonism CI > 1, NAN indeterminate CI) at 50% growth inhibition (Fraction affected). Combinations that are synergistic are colored blue (black otherwise) for each cell lines tested. The inset shows an example of CI calculation for BYL719 and Dasatinib combination in HN12 cell lines based on the corresponding dose matrix (number indicates % cell viability at 48h, n = 3), and Fa-CI curve.

### Experimental testing of predicted DU-SR rescuers via Drug Combinations and siRNA in Head & Neck cancer

In the second experimental validation, we tested the ability of predicted *DU-SRs* to guide new synergistic drug combinations, where the combination of drugs hits both a primary cancer drug target and its predicted *DU* rescuer (Methods). We tested seven such predicted combinational therapies across five different head and neck cancer cell lines. We find that 5 out of 7 combinations predicted are indeed synergistic (**Figure 2b,** Methods, Suppl. Information 4.7, Fig. S6,7). One validated pair involves PI3KCA and mTOR, which are important genes in the PI3K/AKT/mTOR pathway. PIK3CA activates AKT by converting PIP2 to PIP3(*55*), promoting cell growth and survival. mTOR also promotes cell growth and can activate AKT independent of PIK3CA(*56*), thus might compensate for PIK3CA inhibition and explain their synergism.

In a third experiment, we conducted small-scale siRNA experiments to confirm that the synergism observed in above drug combinations is due to direct gene targeting and not due to off-target effects of drugs. To confirm that enhanced sensitivity of the PI3KCA inhibitor, BYL719, is indeed due to mTOR inhibition we directly targeted mTOR by siRNA and showed BYL719 enhanced sensitivity in 4 of these cell lines (Suppl. Information 4.7, Fig. S8). Similarly, we also performed by siRNA experiments to confirm that the enhanced sensitivity of Dasatinib is due to targeting of PIK3CA (Suppl. Information 4.7, Fig. S8).

### Targeting predicted DU-SR rescuers of DNMT1 sensitizes resistant NSCLC cell-lines to DNTM1 inhibitor

In the final and fourth *in vitro* experiment, we tested if targeting predicted *DU* rescuers could sensitize therapy-resistant tumor cells. We picked DNMT1 to test this hypothesis as it is a major hub in the DU-SR network (Figure 1c) and a key oncogene in non-small cell lung cancers (NSCLC). We studied 18 NSCLC cell lines (Methods) that are insensitive to Decitabine (a DNTM1 inhibitor). In each of these cell lines, we pharmacologically inhibited the 13 top predicted DU rescuers of DNTM1. A BLISS(*57-59*) independence model was used to estimate synergism, and its significance was determined by comparing expected vs. observed drug response of drug combinations across all doses tested (Methods). Targeting the predicted rescuers synergistically sensitized these cell lines to Decitabine in 71% of the 234 (13 rescuers x 18 cell lines) conditions tested. In contrast, pharmacologically inhibition of two top predicted DD rescuers of DNMT1 showed the opposite, *antagonistic* effects, in 64% of the 36 conditions tested, with no synergistic effects, as expected (**Figure 3a**). Both the observed synergistic and antagonistic effects across cell lines were significant compared to control drug tested (P < 2.2E-16). We further confirmed the ability of predicted SR interactions to predict resistant tumor sensitization in a large published patient-derived cell line collection(*59*) and mice xenograft (*54*) (Suppl. Information 5.3, 5.4).

**Figure 3.**
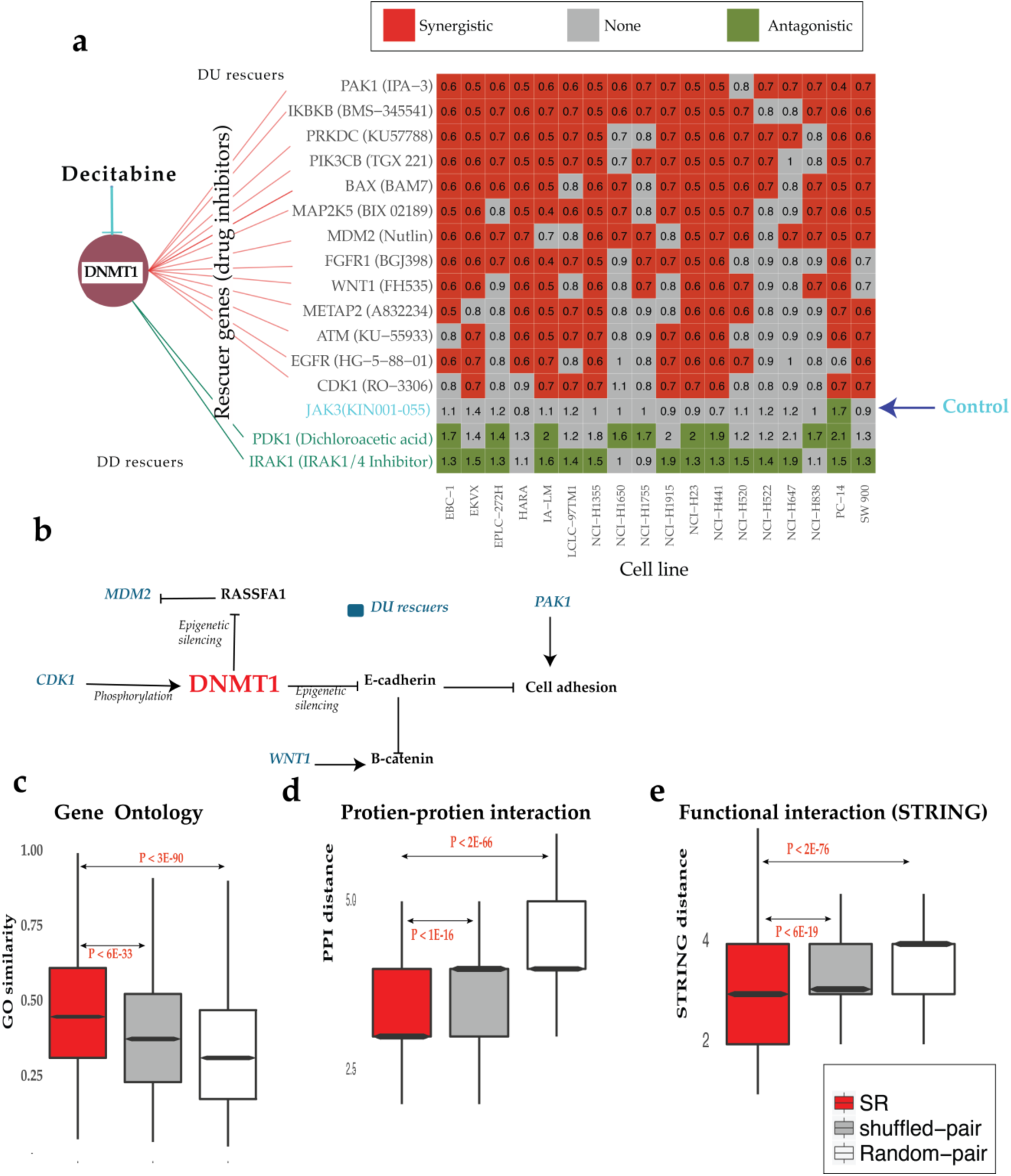
Large-scale experiments testing predicted SRs in NSCLC and studying their functional similarity. (a) Experimental testing of the predicted SR (DU) rescuers of DNMT1 via drug combination experiments in 18 NSCLC cell lines insensitive to Decitabine. The matrix displays drug interactions between Decitabine, a DNMT1 inhibitor, and inhibitors of its predicted rescuer genes (X-axis) across 18 NSCLC cell lines (Y-axis) that are insensitive to Decitabine. Row labels present rescuer genes and their inhibitors. Colors in the matrix show whether the interactions found are significantly synergistic (red), antagonistic (green) or non-significant (in gray). Values in the matrix show average synergism (<1 synergism and >1 antagonism, Methods). 13 predicted SR-DU rescuers (red lines), two predicted DD SR rescuers (green lines) of DNMT1 and one random control (JAK3i) were tested. **(b) Some SR interactions of DNMT1 occur between genes proximally located on the signaling pathway**. DU rescuer genes of DNMT1 are colored blue. **(c-e): Functional similarities between gene pairs in the *DU-SR* network.** Comparison of functional similarities between interactions in (i) the *DU-SR* network (ii) Random-pairs: the network is generated by random pairing between protein-coding genes, having a degree distribution similar to that of the *DU-SR* network (iii) shuffled-pairs: the network is generated by shuffling pairing of the DU-SR network. Functional similarities of genes in each pair were evaluated in terms of their: (c) GO similarity, (d) distance between the paired genes in PPI network, and (e) distances in the STRING network.

The effects of some of the SR interactions validated in drug combination screen described above can be explained by their known biology. E.g.,: (a) First, DNMT1 epigenetically silences E-cadherin(*60*). The silencing results in B-catenin accumulation in nucleus(*61*) that is necessary for maintaining cancer cell stemness. WNT signaling, however, was shown to regulate B-catenin(*62*) independently, explains why WNT1 activation rescues DNMT1 inhibition (**Figure 3b**). (b) Second, DNMT1 also silences RASSFA1, which in turns stabilizes the proto-oncogene MDM2(*63*). Thus, concomitant over-expression of MDM2 could compensate for the loss of RASSFA1 due to DNMT1 inhibition. (c) Third, CDK1 over-expression may compensate DNMT1 inhibition because CDK1 is known stabilize DNMT1 by phosphorylating it(*64*). (d) Finally, PAK1 may compensate for DNMT1 inhibition because it independently regulates cell adhesion and motility. These results testify that some rescue interactions may be explained by molecular interactions between genes proximally located on signaling pathways (*17, 18*) (**Figure 3b**). However, many of the emerging rescue interactions are not, either due to our limited knowledge of signaling pathways or due to functional interactions that go beyond the scope of the signaling pathways.

### Rescuer and vulnerable genes share functional annotations

Our observation that signaling architecture may explain a subset of SR interactions led to the hypothesis that rescuer and vulnerable genes of SR networks may share functional similarities. Several lines of evidence support this hypothesis. First, Gene Ontology annotations (GO) of rescuers of *the DU-SR* network are similar to the GO annotations of their partners (**Figure 3c**). The similarity observed is significantly higher compared to both (i) random network interactions controlled for degree distribution (P < 3E-90), and (ii) shuffled network interactions generated by randomly shuffling the interactions between gene pairs of the DU-SR network (P < 6E-33). Second, *DU-SR* rescuer genes are significantly closer (P < 2E-66 and P < 1E-16 compared to the random network and the shuffled network) to their predicted partners in the protein interaction (PPI) network (**Figure 3d**). Notably, *DU-SR* interactions mediated by direct (physical) protein interactions are enriched in cancer drivers (Fisher exact test P < 6.5E-8, Suppl. Information 3.4). Third, using the STRING database, which integrates multiple resources of direct and indirect associations of protein interactions, we find that partner genes of *DU-SR* network are more likely to be functionally related (**Figure 3e**): Rescuer genes are significantly closer (P < 2E-76 and P < 6E-19 compared to the random network and the shuffled network) to their predicted partner gene in STRING network. Moreover, the observed functional similarities between *DU-SR* pairs are not merely due to co-expression between gene partners; shuffled DU-SR gene pairs with similar co-expression levels as those of predicted DU-SR pairs exhibit significantly less GO similarity (P < 5E-05). An analogous functional similarity was also observed for gene pairs in *DD-SR* network (Suppl. Information 3.8, Fig S2e).

### SR interactions predict drug response in patients

We next turned to evaluate the ability of INCISOR to predict the response of cancer patients to cancer drug treatments(*65, 66*) by analyzing the transcriptomics of their *pre-treated* samples. To this end, we applied INCISOR to identify the rescuers of (the targets of) 28 FDA-approved cancer drugs (for which treatment response data is available in the TCGA collection). During the identification of SR interactions of targets of a given drug, we removed from TCGA patients who were administered with that drug, to remove any potential circularity (Methods, Fig. S10e). To predict the response of an individual patient’s to a given drug, we defined the *drug-tumor SR-score* as the number of upregulated rescuers of the drug’s targets in that patient’s tumor (Methods). We reasoned that a drug is expected to be less effective in tumors where many of its DU rescuers are upregulated. Using a Cox model to control for confounding variables (Methods), we find that the SR-scores predict the patients’ survival after treatment in a statistically significant manner for 22 of the 28 drugs tested (**Figure 4a** shows the result for the 26 drugs tested with hazard ratios > 1, Methods). Evaluating the patients’ response in terms of tumor size (based on the RECIST criteria), we find that the non-responders exhibit significantly higher drug-SR rescue scores than the responders for 14 out of 19 drugs for which tumor size information was available, as expected (**Figure 4b**, Methods). An analysis of independent (non-TCGA) ovarian(*67*) and breast cancer datasets(*68*) further shows that SRs successfully predict both primary and acquired therapy resistance (Suppl. Information 5.2). In contrast, a randomly shuffled network (generated by randomly shuffling rescuer genes for each drug target, maintaining the original SR node degree) fails to predict patients’ response to any of the drugs tested, both in the survival based and response based analyses. Drug-SLs inferred from DAISY(*27*) also showed no predictive signal here (log rank P=0.49). Obviously, INCISOR cannot predict response to drugs without known gene targets.

**Figure 4.**
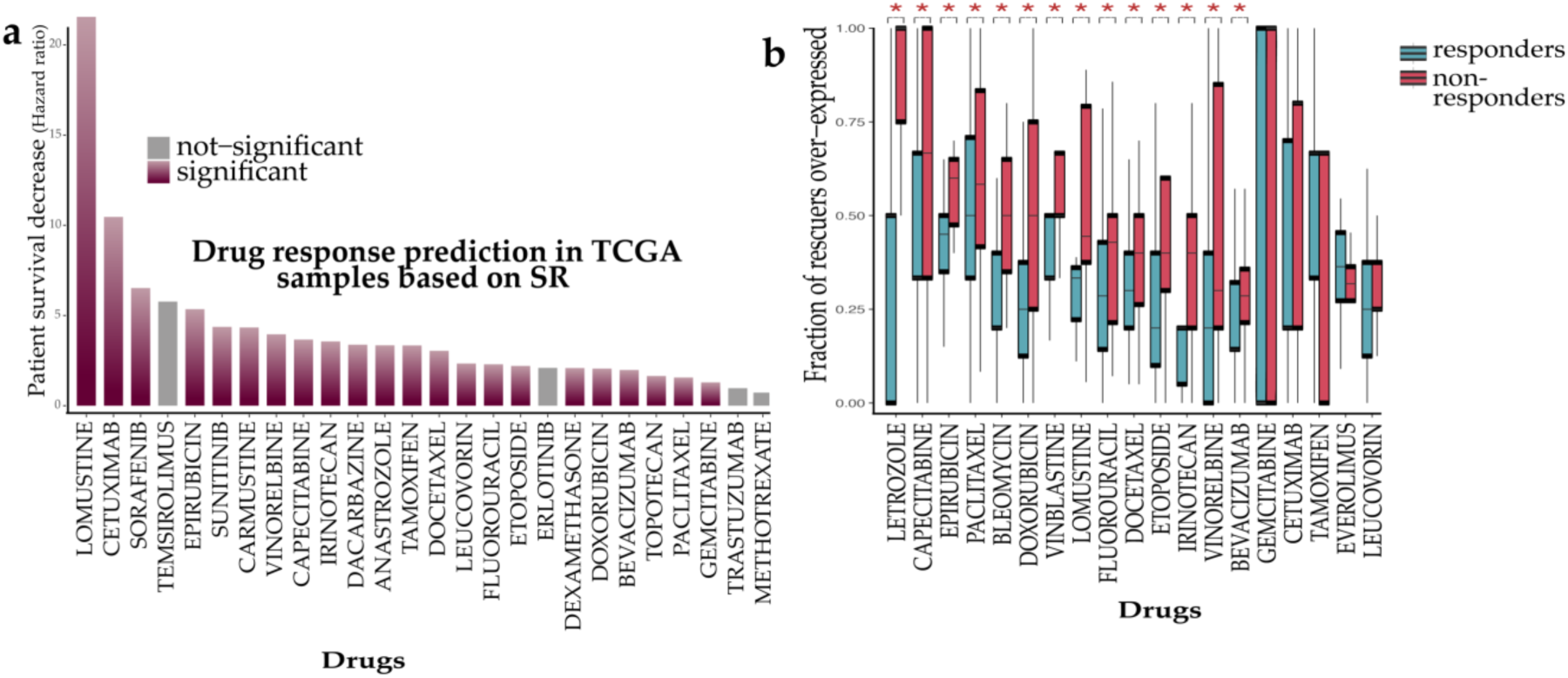
SR networks predict cancer drug response in patients. (a) Prediction of drug response in terms of survival: The Y axis displays the hazard ratio of patients as a function of upregulation of predicted rescuers (Methods). **(b) Analyzing drug response in terms of tumor size reduction (RECIST criteria):** the predicted *DU-SR* rescuers of drugs are differentially over-expressed in non-responding tumors. The Y-axis denotes the fraction of the predicted drug-specific rescuers that are over-expressed (out of all predicted rescuers of that drug) in tumors of responders (red) and non-responders (blue). Significant results are marked by stars (Wilcoxon Rank sum P < 0.05, aggregate Wilcoxon rank sum is P < 2.2E-16, Methods).

### Comparative evaluation of INCISOR’s performance in predicting drug response vs. other recent large-scale genomic methods

We compared the performance of INCISOR with other extant methods for predicting cancer drug response. Iorio et al.(*43*) identified cancer functional events (CFE) and demonstrated that they could be used to predict drug response of 265 drugs *in vitro*. Similarly, Mina et al. (*69*) identified genetic interactions involving these CFEs and demonstrated they predict drug response in cell lines. To systematically evaluate whether these could also determine drug response in patients, we used the occurrence of CFEs and CFE interactions in patients’ tumor as features to build supervised models (Methods) predicting the response for each drug in TCGA. We analyzed 22 FDA-approved cancer drugs in TCGA, including 19 targeted drugs shown in Figure 4b and three drugs without known gene targets (Carboplatin, Cisplatin, and Oxaliplatin). As shown in **Figure 5**, the predicted CFEs of (Iorio et al.) significantly predict drug response for 8, which includes six targeted and two non-targeted therapies drugs. However, the CFE-related genetic interactions identified by Mina et al. do not have a predictive signal for any of these drugs in patients. INCISOR, in turn, outperforms these other methods for 15 out of the 22 FDA drugs analyzed (**Figure 5**). Notably, INCISOR predictions are not based on any supervised training on specific drug response training data and are based on the interactions inferred solely from *pre-treated samples* (supervised predictors of drug response based on INCISOR-predicted rescuers are on average 9.5% more accurate than unsupervised predictors, Fig. S10j).

**Figure 5:**
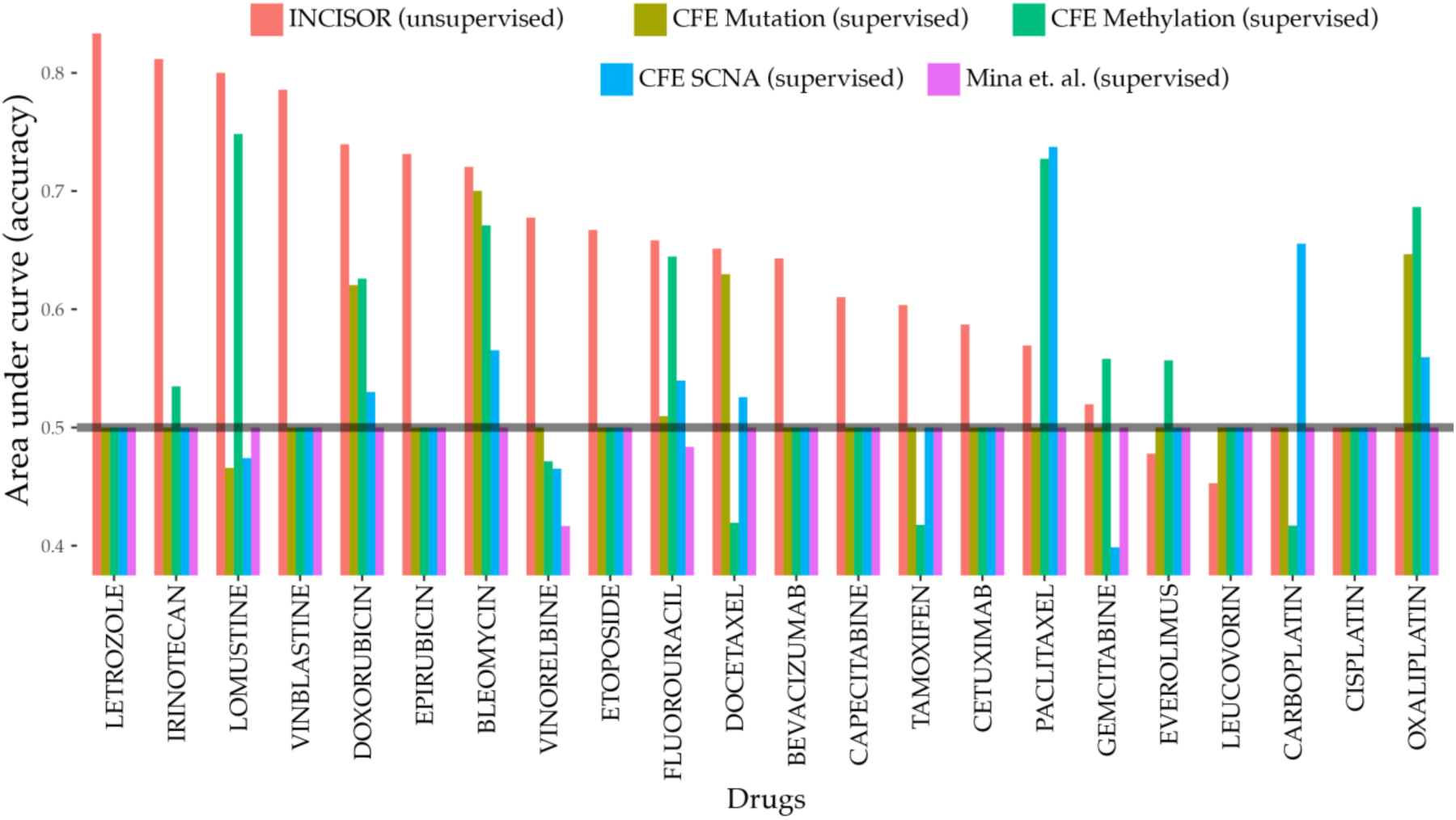
Comparative analysis of INCISOR: A comparative study of INCISOR’s performance (red bars) in predicting patients drug response (TCGA) compared to different CFE based approaches (other colors). The Area under the curve (Y-axis) displays the predictive performance of different methods for 22 FDA approved drugs in TCGA. Predictions of CFE (cancer functional events) identified by Iorio et al. are displayed separately for CFEs inferred from mutation, methylation, and SCNA data.

### SR interactions determine efficacy of immune checkpoint blockades in patients

Finally, we hypothesized that SR-mediated transcriptomic changes mediate resistance to Immune checkpoint blockade (ICB)(*20*). Accordingly, we turned to study the ability of INCISOR to predict SRs that can account for key transcriptomic changes occurring in patients’ tumors following checkpoint immunotherapy and to study the match between the rescuers predicted and key resistance modulators identified in mouse studies. To predict the SR rescuers of the checkpoint genes, we removed *in vitro* essentiality screens (step 1) from the INCISOR pipeline as they are conducted in *in vitro* systems lacking an immune component (Methods). We find that the pretreatment expression levels of INCISOR predicted rescuers of PD1 successfully predict resistance to PD1-blockade in melanoma patients (**Figure 6a**, Prat et al. (*70*) & Hugo et al. (*71*), Suppl. Information 5.5). Similarly, the pretreatment expression of the INCISOR predicted rescuers of CTLA4 successfully predicts patients’ resistance to CTLA4 blockade in melanoma patients (**Figure 6a**, Van Allen et al. (*72*)).

**Figure 6.**
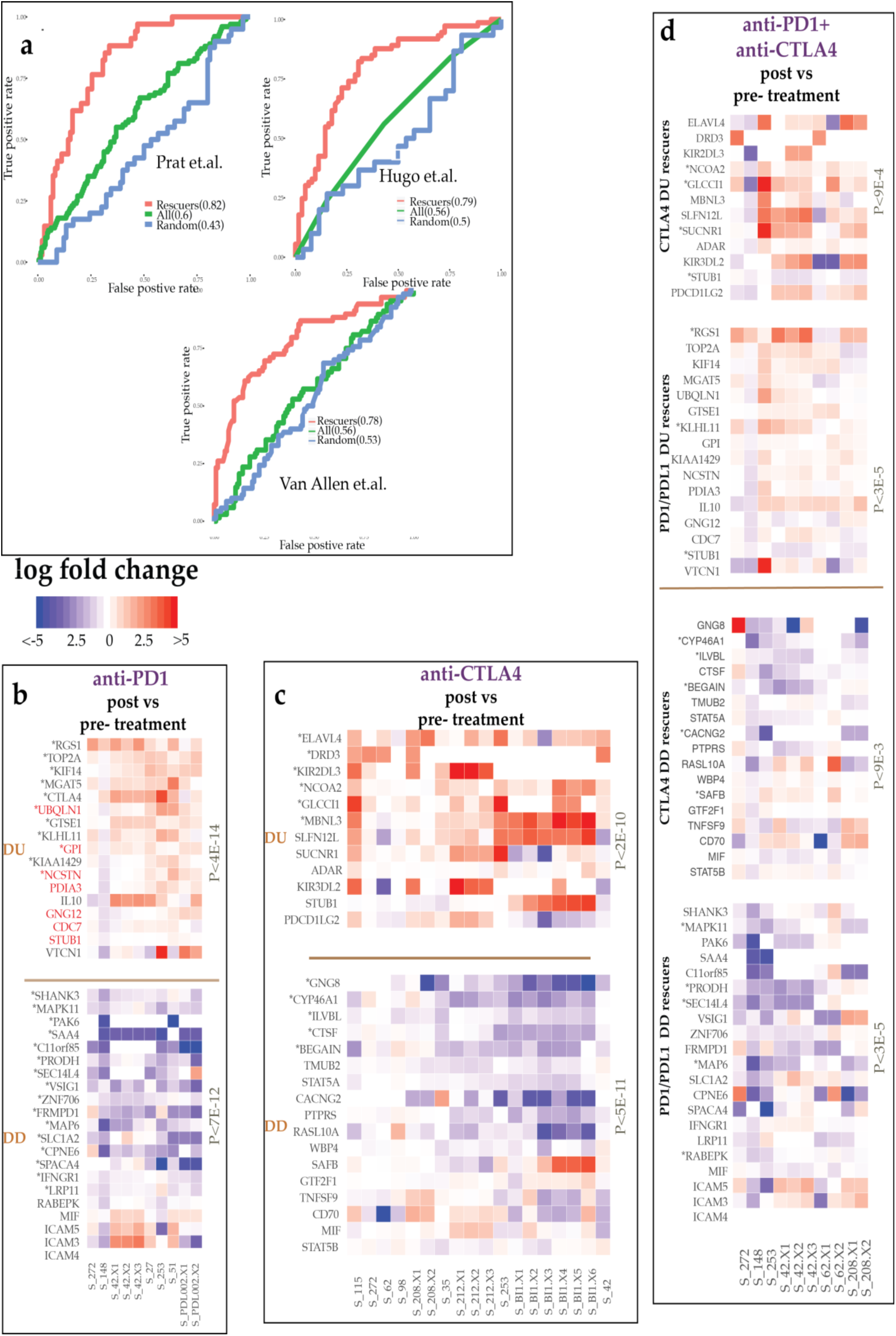
(a) SR predicts resistance to PD1/PDL1 and CTLA4 blockade in patients. Cross-validation accuracy of SR-based supervised predictors in predicting resistance to PD1 (Prat et. al. & Hugo et. al.) and CTLA4 blockade (Van Allen et al.), reported in terms of the corresponding Receiver Operating (ROC) curves. Expression of the predicted rescuer genes of PD1 (CTLA4) was used to train an SVM supervised predictor of PD1 (CTLA4) blockade. For comparison, we also display the ROC curves of supervised predictors trained on the expression of all genes (shown as “All”) and on the expression of genes selected randomly and controlled for the number of rescuers predicted (shown as “Random”). **(b-d) The transcriptomic alterations of rescuer genes post PD1/PDL1 and CTLA4 blockade in patient tumor biopsies:** Their post (vs. pre) treatment expression changes of DU/DD rescuers after anti-PD1 (b), anti-CTLA4 (c), and PD1 + CTLA4 combination therapies (d). Each panel displays the expression fold change of each predicted rescuer gene (rows) for different tumor samples (columns) and the P-value of over-all paired Wilcoxon test of the expression changes observed in paired samples. Significantly altered up/down-regulated genes are marked by (*). Genes marked in red are those whose CRISPR knockdown enhances melanoma sensitivity to anti-PD1 blockade in mice models.

To further study the role of SRs in immunotherapy, we consented 40 patients with metastatic melanoma in ongoing clinical trials for treatment with different ICB therapies and carried out whole transcriptomics profiling of their 90 matched pre-, on- and post-treatment tumor biopsies (Methods, Data available online). 40 biopsies were taken from patients treated with anti-PD1/anti-PDL1 (collated together in the analysis and referred as anti-PD1), 43 biopsies with anti-CTLA4, and 17 with a combination of anti-CTLA4 and anti-PD1 (patients who sequentially underwent from first ICB regiment to another were also considered for individual analysis of the first ICB). Notably, post-treatment biopsies were performed when the patients stopped responding to the ICB, denoting the emergence of resistance.

We find that the top INCISOR predicted DU (DD) rescuers of anti-PD1 therapy (Methods) are preferentially upregulated (downregulated) in anti-PD1 post-treatment tumor biopsies, as expected (paired Wilcoxon p <4E-14) (**Figure 6b** top and bottom panels, their pathway enrichment is provided in Supp. Table S28). Notably, the knockdown of 7 of the 17 predicted upregulated DU rescuers of anti-PD1 therapy has been recently found to promote melanoma’s sensitivity to anti-PD1 blockade in mice models (Hypergeometric enrichment of p < 8E-17; colored red in **Figure 6b**)(*73*). Three of 21 the predicted DD rescuers have also been identified in that study as enhancing resistance, as expected (Hypergeometric enrichment of p < 5.5E-7). More specifically, our results provide evidence in humans that support the mice findings, that gene inactivation of *IFNGR1, RABEPK* and *MIF* induce tumors resistant to PD1 blockade and gene inactivation of *PDIA3, STUB1, CDC7, UBQLN1, NCSTN, GNG12* and *GPI* co-simulates immune response to PD1 blockade in melanoma. Interestingly, we identify *CTLA4* as a DU rescuer of anti-PD1 therapy, supporting the rationale of their combination. Other notable upregulated DU rescuers of PD1 are the immune checkpoint genes *VTCN1* and *TOP2A*. The latter suggest combinations involving DNA topoisomerase inhibitors such as Doxorubicin and Epirubicin. Analogously, top predicted DU (DD) rescuers of anti-CTLA4 therapy were up-regulated (downregulated) in post-treatment tumor biopsies derived from patients treated with anti-CTLA4 therapy ((Paired Wilcoxon p < 5E-11, **Figure 6c;** see Table S29 for pathways enrichment). Notably, we find that anti-CTLA4 blockade can be DU-rescued by a class of inhibitory checkpoints -- Killer-cell immunoglobulin-like receptors (*KIR2DL2* & *KIR3DL3*), which are known to interact with MHC1 and facilitate cell death(*74*), putting forward the potential benefits of combinations targeting these genes. Analyzing samples of post-treatment combination therapy involving both anti-PD1 and anti-CTLA4 we find that many DU/DD rescuers respond as predicted but their individual response is evidently weaker (**Figure 6d**). Notably, the expression of some of the rescuers is altered in the predicted direction already in on-treatment tumor biopsies (Fig. S10h,i), suggesting that these alterations are selected for and thus functionally significant (this does not occur for randomly chosen genes that are not rescuers (P < 2.2E-16)).

## DISCUSSION

In summary, INCISOR prioritizes clinically relevant SRs amongst candidate SRs emerging from *in vitro* screens, by analyzing functional genomic and clinical survival data in an integrated manner. Due to the scarcity of published gold standards of SR interactions, we conducted new large-scale in-vitro experiments to validate our predictions. The paucity of known rescue interactions in the literature further underscores the importance of developing tools like INCISOR. Overall, INCISOR attained precision levels of an average 48% (at 50% recall) in the identification of true SR interaction across all published and new experiments. INCISOR predictions are limited to targeted therapies. Finally, and importantly, we show that SR mediates both primary and adaptive resistance in patients: e.g., we show that the pre-treatment expression data of TCGA tumors is predictive of their response to drug treatments (primary resistance, **Figure 4**) and on the other hand, SRs can predict the post-treatment alterations following checkpoint inhibitors (adaptive resistance, **Figure 6b-d**).

Like many genome-wide approaches, INCISOR has several limitations, including pitfalls arising from gene co-expression and from correlations in the copy number alterations of proximal genes, which may lead to the inference of false positive SRs. We have verified that the SR interactions are not biased towards genes lying on the same chromosome (Suppl. Information 2.3). We aimed to mitigate false positives in the design of INCISOR by selecting candidate SR pairs only when they are additionally supported by the shRNA and phylogenetic data that testify to causal rescue effects. Although INCISOR explains molecular mechanism of resistance to targeted therapies, it fails to capture resistance mechanism of untargeted therapies. Further, for many drugs, resistance can emerge via mechanisms independent of SRs, e.g., resistance due to alteration in drug efflux. Finally, as this is the first genome-wide study of cancer SR interactions, we focused on identifying SRs that are common across many cancer types. Future studies, however, will further identify cancer type-specific SR networks as more data accumulates.

Multiple rescuer genes could rescue and cause resistance to a given cancer drug. Three different strategies could be adapted to prioritize gene target amongst such multiple rescuers to maximize their clinical benefit: First, INCISOR quantifies that extent of the rescue based on its clinical significance observed in patients, which is determined specifically and may be different for each rescuer gene. This could be used for prioritization. Second, post-treatment transcriptomic data from a patient’s tumor, if available, could be used to further narrow down rescuer alterations specific to that tumor. Finally, combining experimental testing in a patient’s tumor material using organoids or PDXs with INCISOR predictions would be a powerful approach to systematically identify true clinically relevant rescuer among the multiple predicted SR rescuers.

This study has focused on the genome-wide prediction of SR interactions. Evidently, different signaling functional and physical interactions may be manifested in these rescue interactions (**Figure 3b-e**). SRs are much less known and studied compared to another type of genetic interactions, known as synthetic lethal (SL) interactions. The difference between SL interactions and *DD-SR* interactions is obvious, by definition. Their difference from *DU-SR* interactions is more intricate: It manifests itself in cells where a given gene is in its wild-type state and its partner interacting gene is knocked down; if the two genes SL-interact there will be no reduction in cellular fitness in that case, but if they DU-SR interact, then the knockdown will reduce cellular fitness (as the rescuer is not up-regulated). Consequently, our results demonstrate that a given cancer drug may be effective in cells where its predicted rescuer is in its wild-type state but may become resistant as it is over-expressed. As expected we found no overlap between predicted DU-SR interactions and SL interactions, predicted by a similar data-mining approach(*30*) (Suppl. Information 2.2).

In summary, we present a novel approach to tackle resistance to targeted and immune cancer therapy by mining thousands of tumors available in TCGA to infer cancer-specific SR interactions. We conducted cell lines experiments demonstrating that targeting predicted DU-SRs could sensitize therapy-resistant tumor cells, identifying synergistic drug combinations. As SR interactions are derived directly from analyzing the patients’ clinical samples, they are more likely to be clinically relevant(*8*) than findings based on cell-screens and mouse models. Our results lay a basis for the development of new combination therapies based on the molecular characteristics of an individual patient’s tumor to proactively overcome resistance in a precision based manner.

## METHODS

### The INCISOR pipeline for identifying SR interactions

INCISOR identifies candidate SR interactions employing four independent statistical screens (Figure 1b), each tailored to test a distinct property of SR pairs. We describe here the identification process for the DU-type SR interactions (Down-Up interactions, where the up-regulation of rescuer genes compensates for the down-regulation of a vulnerable gene (e.g., by an inhibitor compound, Fig. S1a). Then we discuss how to modify DU-INCISOR to detect the other SR types (DD, UD, and UU). We identify pan-cancer SRs (that are common across many cancer types) analyzing gene expression, somatic copy number alteration (SCNA), and patient survival data of TCGA (*75*) from 8,749 patients in 28 different cancer types. INCISOR also integrates predictions from TCGA data with genome-wide shRNA(*39, 40, 42*) and drug response (*43, 44*) screens in around 720 cell lines composing in the total of 2.3 million shRNA measurements. The same approach can be used to identify cancer type specific SRs, in an analogous manner. INCISOR is composed of four sequential steps (an FDR threshold was set 0.05 for each step):

(1) *In-vitro screening (using in vitro cancer data):* Mining large scale in vitro shRNA and drug response datasets, INCISOR examines all possible gene pairs to identify putative SR. The screen adopts an analogous approach (*29*) to mine shRNA screen in a reference collection of cell line to identify pairs where vulnerable genes V and rescuer genes R fulfill the following two conditions: (i) knockdown of the V exhibits an increase in cell growth in cell lines with R upregulated (relative to cell line with R downregulated), and (ii) knockdown of the R is lethal in cell lines where V downregulated. We use both gene expression and SCNA data to identify such putative SR as follows: We divide the cell lines into those having high or low expression of gene R and compare the cell growth of V in these two the groups using a Wilcoxon rank sum test. Similarly, we determine conditional essentiality of R in two groups with high and low expression of R. SCNA based conditional essentiality is determined analogously, where the groups are determined based on SCNA of V or R. Pairs that pass both tests (i) and (ii) either using gene expression or SCNA are referred as putative SR.

Analogously, Drug-response screen is based on the condition that treatment of a drug inhibitor targeting vulnerable gene V will be resistant in cell lines where its rescuer gene R is active. Accordingly, we integrate gene-expression and SCNA of cell lines with drug response to identify such putative SR pairs that exhibit significant *in vitro* conditional resistance. Putative SR pairs **significant either in shRNA screen or Drug-response screen** following the standard FDR correction (*76*) are passed on to the next screen.

(2) *Molecular survival of the fittest (SoF, analyzing tumor molecular data)*: This screen mines gene expression and SCNA data of the input tumor samples to identify vulnerable gene (V) and rescuer gene (R) pairs having the property that tumor samples in the *non-rescued* state (that is samples with underactive gene V and non-overactive gene R, activity states 1 and 2 in Fig. S1a) are significantly less frequent than expected, whereas samples in the *rescued* state (that is samples with under-active gene V but over-active gene R) appear significantly more than anticipated (testifying to the positive selection of *rescued* state of the pairwise interaction). The significance of the enrichment/depletion of rescued/non-rescued state is determined via a hypergeometric test followed by standard false discovery rate correction. A gene is defined as inactive (respectively, overactive) if its expression level is less (greater) than the 33rd-percentile (67th-percentile) across samples for each cancer type (to control for cancer-type). Otherwise, it is considered to have a normal activation level. Out of total N tumor samples, if n1 (n2) is the number of samples in the rescued/non-rescued state using specific activation level of gene R (V) independently, k is the number of samples in the activity state using both genes R and V, the significance of enrichment/depletion of the observed number of samples in the rescued/non-rescued state is determined using hypergeometric test: *hypergeomatric(k, n1, N, n2)*. Enrichment/depletion of the activity state using SCNA is set analogously. Pairs significant (FDR < 0.05) in both SCNA and mRNA are passed on to the next screen.
(3) *Clinical screening (using patient survival data):* This step selects a gene pair as SR if it has the property that tumor samples in *rescued* state (that is samples with underactive gene V and overactive gene R) exhibits significantly poorer patient’s survival and samples in *non-rescued* state tumors exhibits better survival than rest of the other samples. Specifically, INCISOR uses a stratified Cox proportional hazard model to check such observed associations of SR *rescued*/*non-rescued* state are significantly larger compared to the expected additive survival effect of their individual genes, while controlling for confounding factors including cancer type, sex, age, genomic instability, tumor purity, and race (shown here for expression analysis for an activity state *A* and a similar model is used to analyze SCNA data):

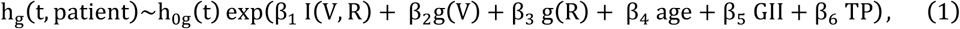

Where g is a variable over all possible combinations of patients’ stratifications based on cancer-type, race, and sex. h_g_ is the hazard function (defined as the risk of death of patients per unit time), and h_0g_(t) is the baseline-hazard function at time t of the gth stratification. The model contains five covariates: (i) I (V, R) : indicator variable representing if the patient’s tumor is in the activity state *A*, (ii) g(V) and (iii) g(R): gene expression of V and R, (iv) age: age of the patient, (v) GII: genomic instability index of the patient, and (vi) TP: tumor purity. The βs are the unknown regression coefficient parameters of the covariates, which quantify the effect of covariates on the survival. All covariates are quantile normalized to N(0,1). The βs are determined by standard likelihood maximization (*77, 78*) of the model using the R-package “Survival.” The significance of β_1_, which is the coefficient for the SR interaction term is determined by comparing the likelihood of the model with the NULL model without the interaction indicator I (A, B) followed by a likelihood ratio test and Wald’s test (*77, 78*), i.e.,

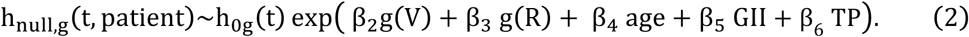 The p-values obtained are corrected for multiple hypothesis testing. We pass a putative SR pair to the next screen if its *rescued state* exhibits significantly poorer survival and the *non-rescued state* exhibits better survival both regarding mRNA and SCNA (all FDR < 0.05).

Tumor purity is obtained for each TCGA samples from Aran et al. (*45*). They combined following four methods to estimate an aggregate estimate of tumor purity: (i) ESTIMATE(*79*), (ii) ABSOLUTE(*80*), (iii) LUMP; and (iv) IHC. We control for tumor purity estimated from each of these four methods in addition to the aggregated tumor purity in the survival analysis.

1. (4) *Phylogenetic profiling screening:* we further filter and select SR pairs composed of genes having high phylogenetic similarity, motivated by the findings of Srivas et al. (*28*). This is done by comparing the phylogenetic profiles of the SR-paired genes across a diverse set of 87 divergent eukaryotic species adopting the method of Tabach et al. (*81, 82*). The resulting matrix of the phylogenetic scores of all candidate genes is clustered using a non-negative matrix factorization (NMF) (*83*), and the Euclidian distance between the cluster membership pattern of each gene in given candidate pair is computed. The significant (empirical-FDR < 0.05) phylogenetically similar pairs are predicted as the final set of SR pairs.

Both SCNA and transcriptomic data are useful to determine the SR interactions because gene expression is closer to the phenotype than SCNA, and the later are inherited during clonal cancer growth. Therefore, INCISOR (SoF, Clinical and shRNA screenings) uses both SCNA and gene expression as independent evidence of gene-activity to infer SR interactions that are more likely to be causal. Specifically, we check that the statistical tests used in each screening are significant both in terms of SCNA and gene expression for an SR interaction to be significant.

To process half a billion gene pairs for around 9,000 patient tumor samples in a reasonable time, the most computationally intensive parts of INCISOR are coded in C++ and ported to R. Further; INCISOR uses open Multiprocessing (OpenMP) programming in C++ to use multiprocessor in large clusters. Also, INCISOR performs coarse-grained parallelization using R-packages “parallel” and “foreach”. Finally, INCISOR uses Terascale Open-source Resource and QUEue Manager (TORQUE) to uses more than 1000 cores in the large cluster to efficiently infer genome-wide SR interactions.

#### Applying INCISOR to construct the DD-SR network

##### Constructing the DD-SR network

We modified INCISOR in DD-SR network inference to account for the fact that rescuer gene down-regulation leads to synthetic rescues. In DD, the *rescued* state is defined as co-inactivation of vulnerable (V) and rescuer gene (R); and *non-rescued* state is defined as underactive gene V and active gene R (Fig. S1b). Accordingly, the four screens of INCISOR, described above for DU identification, were modified as follows: (i) **SoF** and **Survival screening:** The statistical tests (i.e., hypergeometric test and Cox regression) are modified so as to account for DD interactions that have different activity states (i.e., *rescued* and *not-rescued* states, Fig. S1b). (ii) **shRNA screening**: Similarly, the *conditional* knockdown of a DD rescuer gene, now increases the cell proliferation due to activation of DD synthetic rescue. The significance of the increase in the cell proliferation due to a rescuer down-regulation is quantified in an analogous manner using Wilcoxon rank sum test. (iii) **Phylogenetic screen**: it remains the same as the case of DU identification (refer to Suppl. Information 2 for additional details).

#### Interactive SR networks

The four types of SR networks for pan-cancer were created using Cytoscape (*84*) and are accessible online in an interactive manner at http://www.umiacs.umd.edu/~vinash85/private/SR/(with username: “sr” and password: “sr123”).

#### Genomic instability index

Genomic instability index measures the relative amplification or deletion of genes in a tumor based on the SCNA. Given s_*i*_ be the absolute of log ratio of SCNA of gene *i* in a sample relative to normal control, GII of the sample is given as (*85*):

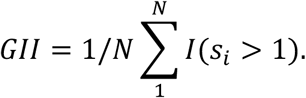

### Calculation of INCISOR interaction-score

INCISOR evaluates each of the candidate SR gene pairs based on the strength of their SR interactions. We define *INCISOR interaction-score,* which combines the significance levels of the four statistical tests in the INCISOR pipeline. First, for each screen, the statistical significance levels of all gene pairs tested were rank-normalized to a value between 0 and 1 (with 0 representing a pair with the highest significance and 1 with the lowest). The final INCISOR interaction-score for a gene pair *i* is given as:

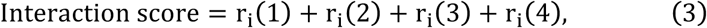

where r_i_(k) represents the rank normalized value of the *k*^th^ screen of INCISOR.

### Mapping of drugs to their gene targets

The drugs were mapped to their targets based on the mapping reported in CCLE, CTRP, and DrugBank (*43, 86, 87*) with exception of target genes whose mechanism of action is explicitly denoted as an agonist in DrugBank.

### Effect size via Cohen’s d

Throughout the manuscript, whenever applicable, to quantify a difference between two groups, we use an effect-size measure called *cohen’s d* (*88*). It is defined as the difference of means divided by pooled standard deviation. Given s_1_ and s_2_ as standard deviations of two groups and n_1_ and n_2_ are a number of samples in each group, the pooled standard deviation is defined as:

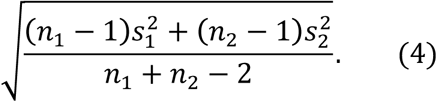

### Pathway enrichment

GO, and KEGG enrichment analyses were conducted using R-packages *clusterprofiler* and *GOFunction* using default settings.

### Precision and Recall

Using standard definitions, we define INCISOR’s precision as the fraction of true SR interaction among the predicted SR interaction by INCISOR. The INCISOR’s recall is defined as the fraction of true SR interactions that are retrieved by INCISOR among all true SR interactions.

### Benchmarking DU-SR networks using literature compiled SR interactions

The seven datasets of the published SR interactions were compiled using extensive literature survey of large clinical and experimental studies (Datasets, dataset pairs, and associated publications are listed in Table S9). Each dataset consists of a drug and experientially and/or clinically validated genes whose over-expression cause resistance to the drug treatment in patient samples/cell lines. In each study, the pairings between the drug targets (vulnerable genes) and the corresponding resistance-causing genes (rescuer genes) form the positive set; and the pairings between the targets and all other genes tested, which do not exhibit resistance, forms the negative set. Using the INCISOR *interaction-score* of individual SR-pairs as the prediction for the strength of SR interaction, we performed standard ROC and precision-recall analysis (Suppl. Information 4.1).

### Constructing the drug DU-SR network

To remove any potential circularity in drug response prediction, for each drug analyzed, we excluded from TCGA dataset the samples of the patients who were treated with that drug. Next, we applied INCISOR to the remaining TCGA samples to identify rescuers of the targets of the drug. The resultant drug-DU-SR network applied for 28 targeted drugs constitutes 182 rescuer genes of 24 drug targets (Suppl. Information 5.1).

### Predicting pan-cancer drug response in patients

#### Prediction of drug response using patient survival

Using the drug-DU-SR network, we analyzed 4,328 TCGA samples, which is the collection of samples of patients who were treated with the drugs that were administered to at least 30 patients in TCGA. We predicted that patient tumors would be resistant to drug treatment if multiple DU rescuer genes of the drug targets are upregulated in their tumor. Therefore, the number of rescuer gene over-expressed will be predictive of patients’ drug response. Accordingly, for a drug tested **D** and each patient administering **D**, we estimate the fraction (***C***) of DU rescuer genes upregulated of its drug targets (deduced from their gene expression and SCNA values in the pre-treatment tumor sample) in the patient sample. To predict the response of TCGA patients treated, we evaluated the association of ***C*** with the patients’ survival using stratified Cox model, which also controls for confounding factors(cancer-type, age, sex, and race) as follows:

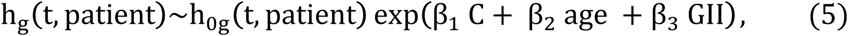

Where h_g_, h_0g_, βs *age,* and GII are defined as in the equation (**1**). Co-variates ***C**, age,* and GII are quantiles normalized to N(0,1). The significance of β_1_, which is the coefficient of ***C**,* is determined by comparing the likelihood of the model with the NULL Cox model, which is similar to **(3)** but without the covariates ***C**,* followed by likelihood ratio and Wald’s tests (*77, 78*). As evident, SRs can be successfully used to predict drug response in an unsupervised manner (which is hence less prone to over-fitting).

#### Prediction of patient drug response based on post-treatment patient tumor size

We evaluated the performance of our prediction vs. TCGA drug response based on patient tumor size following the treatment. Based on RECIST drug response profile of 3872 patients in TCGA, which were annotated into Complete Response (CR), Partial Response (PR), Stable Disease (SD), and Progressive Disease (PD), we divided the samples into responders (CR and PR) vs. non-responders (PD and SD). To determine the ability of SR to predict drug response of each drug, we compared the fraction of the DU rescuers (of the drug’s targets) upregulated in patients’ tumors (***C***), and their significance is determined using Wilcoxon rank sum test.

### Comparative performance of INCISOR in predicting drug response

#### DU-SR based unsupervised predictor

To predict the response of a drug in an unsupervised manner, we first identified responders and non-responders in TCGA dataset. The fraction of over-expressed rescuers of targets of the drug in each patient was used to estimate the area under the curve (AUC). If AUC > 0.5 and mean fraction of over-expressed rescuers was higher in responders compared to non-responders (1-AUC) was used as the final estimate of AUC.

#### Supervised prediction of patient response using CFE (*43*)

The list of CFEs were collected from Iorio et al. (*43*). It provides four distinct types of CFEs: (i) mutation, (ii) methylation, and (iii) somatic copy number alteration (SCNA). Predictive performance each type of CFEs were evaluated individually. Using TCGA data, we generated a matrix of CFE occurrence across all TCGA patients. The CFE occurrence matrix was used as features to train supervised models for predicting patients’ response for each of 22 FDA approved drugs as follows.

To predict the response of a drug in a supervised manner, we first identified responders and non-responders in TCGA dataset. Given the CFE occurrence matrix as features described above, we built a random forest based supervised predictor that discriminates responders from non-responders. The random forest was preferred over SVM because its performance was superior as compared to SVM for this prediction task. Twofold cross-validation was used to estimate AUC.

#### Supervised prediction of patient response using CFE interaction (Mina et al. (*69*))

The list of CFEs were collected from Mina et al. (*69*). ANOVA p-value < 0.05 was used to filter out non-significant drug and CFE pairs, resulting 1444 drug CFE pairs with significant association. CFE occurrence in TCGA patients was downloaded from ciriellolab.org/select/select.html, and was used to identify if CFE-pairs co-occur in the patient’s tumor, which is represented as an indicator variable. To predict the response of a drug, we used CFE-pairs reported to significantly associated with the drug by Mina et al. as features. The corresponding matrix of CFE-pairs co-occurrence in patients was used to train a supervised model in an analogous manner described above for Iorio et. al.

### Experimental testing of INCISOR-predicted SR interactions involving mTOR

We used Rapamycin because it is a highly specific mTOR inhibitor and hence enables targeting of a predicted rescuer gene by a highly specific drug, combined with the ability to knock down predicted vulnerable genes in a clinically-relevant lab setting. Rapamycin is known to specifically targets mTOR in its complex 1 (*89*). Its selectivity stems from the need to act on a protein FKBP12, which binds to the FKBP12-binding region (FRB) in mTOR (*90*). This was confirmed in our earlier work (*91*) (particularly in the HN12 cell line, which we used in our experiment) by expressing an FRB mutant mTOR that cannot bind to the rapamycin-FKBP12 complex, which rescued these cells from the anti-tumor effect of Rapamycin in vitro and in vivo. This retro-inhibition approach further supported the specificity of rapamycin for mTOR, in a biologically relevant context. And therefore we choose HN12 to conduct this experiment.

Fig. S5a provides the overview of overall experimental procedure (Suppl. Information 4.6). We performed the shRNA knockout and mTOR inhibition in the following steps. 2214 gene kinases (Table S10) were knocked down in HN12 cell lines, after which mTOR was inactivated via Rapamycin treatment. HN12 cells were infected with a library of retroviral barcoded shRNAs at a representation of ~1,000 and a multiplicity of infection (MOI) of ~1, including at least two independent shRNAs for each gene of interest and controls. At day three post infection cells were selected with puromycin for three days (1µg/ml) to remove the minority of uninfected cells. After that, cells were expanded in culture for three days, and then an initial population-doubling 0 (PD0) sample was taken. For in vitro testing, the cells were divided into six populations, three were kept as a control, and three were treated with Rapamycin (100nM). Cells were propagated in the presence or not of a drug for an additional 12 doublings before the final, PD13 sample was taken. shRNA barcode was PCR-recovered from genomic samples and samples sequenced to calculate the abundance of the different shRNA probes. From these shRNA experiments, we obtained cell counts for each gene knock-down at the following three time points: (a) post shRNA infection (PD0, referred as initial count), (b) shRNA treatment followed by either Rapamycin treatment (PD13, referred as treated count, 3 replicates) or control (PD13, referred as untreated count, 3 replicates) (c) shRNA infected cell injected to mice (tumor, referred as in-vivo count, 2 replicates).

Significant experimental (DD) rescue event was determined by using Mageck (*92*), where read counts from pooled shRNA were the first median normalized for both rapamycin treated samples (3 replicates) and controls (3 replicates). The difference of read counts in rapamycin-treated samples and controls was then modeled as a negative binomial model and was used to test if the difference is significant for each gene tested in pooled shRNA (*92*).

Next, we applied INCISOR to TCGA to specifically predict DD-SR interactions between 2214 genes tested and mTOR as DD rescuer. Each gene pairing (between 2214 genes and mTOR) were quantified using INCISOR interaction-score. The score was used to estimate precision and recall.

To quantify the lethality of vulnerable knockdown in the experiment, we performed a one-sided Wilcoxon rank-sum test between initial normalized count with in vivo normalized count for in vivo lethality (and with the untreated normalized count for in vitro lethality). To compare rescue effect of Rapamycin treatment between shRNA knockdown of mTOR’s vulnerable gene partner and control gene knockdown, we performed a one-sided Wilcoxon rank-sum test between Rapamycin effects of mTOR partner vulnerable genes and control genes.

Fig. S5b was generated as follows: To obtain normalized counts at each time point, cell counts of each shRNA at each time point were divided by corresponding a total number of cell count. To estimate cell growth rate at treated, untreated and in vivo time points for each gene X, normalized counts were divided by initial normalized count as follow:

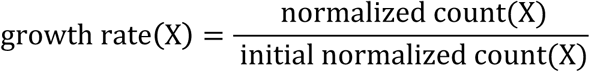

Effect of Rapamycin treatment on cell growth on knockdown of gene X was calculated as:

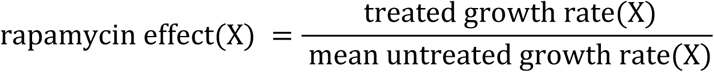

### Experimental Testing of SR-predicted Synergistic Drug Combinations in head and neck cancer cell lines

Seven drug combinations tested in this experiment were chosen as follows. Among the important cancer genes captured by DU-SR network, we focused on testing SR interactions between 5 important HNSC oncogenes (mTOR, PIK3CA, KIT, AKT, and PTK2). INCISOR predicted 5 SR-DU interaction between these oncogenes. Two different inhibitors (Rapamycin and ink128) were included for mTOR, and one inhibitor each for PIK3CA, KIT/SRC, AKT and PTK2 were included in the experiment. This resulted seven combinations tested (Figure 2b).

Rapamycin was purchased from LC Laboratories (Woburn, MA). Dasatinib, Erlotinib, BYL719, INK128 were purchased from Selleckchem (Houston, TX).Vita-Blue Cell Viability Reagent was purchased from Bio-tools (Jupiter, FL). CAL33, HN12, Detroit 562 and SCC47 cell lines were cultured in 96-well-plate, then treated with drugs for 48 hours (Raw Data in Table S11-17). Assays were performed according to the manufacturer’s instructions. Combination index for quantitation of drug synergy was analyzed by CompuSyn software (*93, 94*). CI values represent synergism (CI<1), additivity (CI=1), and antagonism (CI>1), respectively (Suppl. Information 4.7).

### Experimental Testing of SR-predicted rescuers via siRNA

siRNAs for non-targeting control and PIK3CA were purchased from GE Healthcare (two ON-TARGETplus PIK3CA siRNAs 5’-GCGAAAUUCUCACACUAUU, and 5’- GACCCUAGCCUUAGAUAAA, Lafayette, CO). siRNAs for mTOR were purchased from Sigma (two MISSION^®^ siRNA human mTOR SASI_Hs02_00338641 and SASI_Hs01_00203144). Cells were cultured in 96-well-plate, transfected with lipofectamin RNAiMAX reagent (Life Technologies, Carlsbad, CA) for 24 hours, then cells were treated with drugs for another 48 hours (Raw Data in Table S18,19). Viability assays were completed as previous described (Suppl. Information 4.7).

To test whether the KD of INCISOR-predicted rescuers acts as sensitizers, we checked the sensitivity of cells to the primary drug increases upon the KD of the rescuer. The efficacy of the rescuer KD to sensitizes cancer cells to a primary drug was estimated as the percentage increase in the sensitivity to the drug following the rescuer KD relative to the sensitivity of the primary drug alone without rescuer KD. Specifically, we used a targeted siRNA to knockdown the specific rescuer gene, while an untargeted non-specific siRNA was used as a control. Cell counts were measured for the untargeted/targeted siRNA post primary drug treatments. The normalized-response of (targeted/untargeted) siRNA-treated-cells to drug treatment was quantified as a change in cell counts relative to the cell counts following the respective siRNA inhibition alone. Next, using this normalized-response, DRC was estimated for both targeted and untargeted siRNA using DRC R-package. Percentage increase in sensitivity of the primary therapy (Y-axis, Fig. S8i) due to the rescuer siRNA-KD was estimated as the percentage decrease in IC50 of the combination of primary drug treatment and siRNA inhibition relative to the primary therapy in untargeted siRNA combination. The significance of the increase in drug response was estimated using a standard ANOVA test.

### Drug combination testing of SR interaction involving DNMT1

#### Drug combination Screen

Drug screening was performed using automated liquid handling in a 1536 well plate format(*59*). The drug doses used were chosen based on previous single agent screening at the Center for Molecular Therapeutics of the Massachusetts General Hospital Center for Cancer Research (Table S24).

The screen of two drug A and B was performed in a 1×5 format with one dose of drug A combined to 5 doses of drug B and compared to the effects of the five doses of drug B alone. The five doses of drug B followed a four-fold dilution series (Table S24, 25).

Cells were seeded at densities optimized for proliferation based on the pre-screen experimental determination in 1536 well plate format. Cells were seeded, placed overnight at 37°C and drugs added the next day using a pin tool. After five days in drug cells were fixed permeabilized and nuclei stained in a single step by adding a PBS Triton X100 / Formaldehyde / Hoechst-33342 solution directly to the culture medium. Final concentrations: 0.05% Triton X-100 / 1% Formaldehyde / 1 ug/ml Hoechst-33342. Plates were covered and placed at 4^°^C until imaging.

Imaging was performed on an ImageXpress Micro XL (Molecular Devices) using a 4X objective. Cell nuclei enumeration was performed using the MetaXpress software, and count accuracy were routinely checked visually during acquisition. The screening was conducted in two replicate (two separate 1536 well plates, Table S25).

#### Calculation of drug combination synergy score

Due to a limited number of dose combination used in the experiments per each drug pair (1 concentration of Decitabine (five replicates) and five concentration of each rescuer inhibitor (2 replicates)), Fa-CI analysis is not feasible. The drug dose tested are provided in Table S24.

We used Bliss independence model (*57-59*) to determine synergistic drug combination which is suitable per such experimental setting. More specifically, to determine Decitabine synergism with a rescuer inhibitor (R) tested in a cell line, we compared following two ratios of experimentally determined cell counts for each dose (C) of rescuer inhibitor (Table S25):

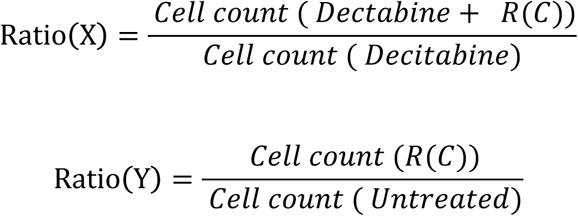

Where *Cell count (X)*, denotes cell count following the treatment of X. R(C) denotes rescuer inhibitor at dose C. The effect size of synergism at dose C of rescuer inhibitor was estimated as 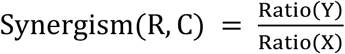. This calculation is separately done for each of two replicates of R dose, generating 10 data points (5 dose x 2 replicate) for each rescuer inhibitors.

The final synergism of rescuer inhibitor R was estimated as the median of Synergism of the 10 data points, i.e., 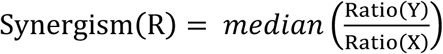. Significance of synergism of R was estimated by a Wilcoxon rank sum test comparing Ratio(Y) and Ratio(X) of the 10 data points of R. Finally, R was estimated to be significantly synergistic with Decitabine, if Synergism R > 1.25 and p-value adjusted for multiple hypothesis corrections is less .05 (i.e., FDR < 0.05, Table S26).

Analogously, R was considered to be significantly antagonistic with Decitabine, if Synergism R < 0.75 and p-value adjusted for multiple hypothesis corrections are less .05 (i.e., FDR < 0.05, Table S26).

### Functional similarity of SR

#### Gene ontology similarity

Gene ontology semantic similarity(*95*) was used to quantify the similarity of GO terms between a gene pair. When multiple GO terms were associated with a gene, similarity between all combinations of rescuer GO terms, and vulnerable GO terms were calculated, and the maximum of these scores was taken as final similarity score (average of scores as final similarity gives similar result qualitatively). Distribution of GO similarity of DU-SR pairs was compared with two sets of controls: (a) shuffled network: interactions between rescuer and vulnerable genes of DU-SR network randomly shuffled, and (b) random network: gene pairs selected randomly from all protein-coding genes and controlled for similar degree distribution as the original DU-SR network. For each set of control, we determined the similarity measure in an analogous manner as described above for DU-SR network. Wilcoxon rank sum test was used to calculate the significance of GO similarity of DU-SR network relative to each control.

#### PPI distance

IGraph was used to estimate the distance between two genes in human protein-protein interactions (PPI) network compiled from (*96, 97*). The PPI distance between gene pairs was compared with two controls the random network and shuffled network as described above.

#### STRING database distance

The STRING network version 10 was downloaded using R-package StringDB. STRING database is composed of gene pairs that are likely to share functional similarities. The functional similarity scores provided in the database were estimated using various sources including direct (physical) and indirect (functional) associations. The comparison control networks were made analogous manner as in case of GO similarity described above.

For DD-SR network GO similarity, PPI distance, and STRING database distance were estimated analogously (Suppl. Information 3.8).

#### PPI specific DU-SR interactions

To identify DU-SR interactions likely to be mediated by PPI interactions, we applied INCISOR on human PPI network compiled from (*96, 97*). The details of the analysis and resultant network are provided in Suppl. Information 3.4. The enrichment of cancer driver genes in PPI-SR network was calculated using Fisher exact test.

### Identification of DU and DD rescuers of immune checkpoints

To identify rescuers of immune checkpoints we removed filtering step 1 (*in vitro* essentiality screens) from INCISOR because the dataset used in step 1 were conducted in *in vitro* models that are deficient of the immune system. Next, we applied the INCISOR to the TCGA to identify the DU and DD rescuers of PD1/PDL1 and CTLA4 down-regulation. We call an SR identified by INCISOR as clinically significant if the interaction shows association with survival either in pancancer or melanoma cohort in TCGA patient dataset.

### Immunotherapy samples patient samples collection and processing

#### Patient samples

A cohort of patients with metastatic melanoma treated was enrolled in clinical trials ongoing at Massachusetts General Hospital for treatment with three immune checkpoint blockades: (i) anti-PD1 or anti-PDL1 (collated together in the analysis and referred as anti-PD1), (ii) anti-CTLA-4, and (iii) combination of anti-PD1 and anti-CTLA-4. Patients were consented for tissue acquisition per Institutional Review Board (IRB)-approved protocol (*98*). These studies were conducted according to the Declaration of Helsinki following informed consent (DF/HCC protocol 11- 181) was obtained from all patients.

#### RNA sequencing (RNA-seq)

Tumors were biopsied or surgically removed from the consented patients and snap frozen in liquid nitrogen or fixed in formalin. Qiagen AllPrep DNA/RNA mini or AllPrep DNA/RNA FFPE kit was used to purify RNA from the frozen or fixed tumor biopsies. RNA libraries were prepared from 250 ng RNA per sample using standard Illumina protocols. Samples were treated with ribo-zero, and then Epicentre’s ScriptSeq Complete Gold kit was used for library preparation. The quality check was done on the Bioanalyzer using the High Sensitivity DNA kit, and quantification was carried out using KAPA Quantification kit. RNA sequencing was performed at Broad Institute (Illumina HiSeq2000) and The Wistar Institute (Illumina NextSeq 500). BAM files of raw RNA-seq data were used to summarize read counts by featureCounts (*99*) with parameters that only paired-ended, not chimeric and well mapped (mapping quality >= 20) reads are counted (Data available online).

Differential expression analysis was conducted by generalized linear model implementation (*100*) of R package “edgeR” and following a standard pre-processing of read count analysis (*101*). Transcript per million (TPM) were used to estimate fold change.

If a patient is treated sequentially with ICBs A and B and the biopsies are available pretreatment, post-A-treatment and post A+B treatment. Comparison of pretreatment vs. post-A-treatment of the patient was considered in the analysis for the resistance of therapy A. In case of multiple biopsies for pre-, on- or post-treatment are available per patient all biopsies were considered in the analysis as follows. For the analysis of differential expression using edgeR, an indicator variable per patient was introduced in design matrix as recommended in reference manual of edgeR, which controls for individual specific transcriptome. To calculate the fold change displayed in **Figure 6**, mean of TPM was taken in case of multiple pre-treatment biopsies per patients; and in case of multiple post- or on-treatment each biopsy were displayed in the figure (subscripted by “.X[biopsises number]”).

